# Transformer-Based Deep Learning Model with Latent Space Regularization for CRISPR-Cas Protein Sequence Classification

**DOI:** 10.1101/2024.03.02.583136

**Authors:** Bharani Nammi, Sita Sirisha Madugula, Pranav Pujar, Vindi Mahesha Jayasinghe Arachchige, Jin Liu, Shouyi Wang

## Abstract

The discovery of the CRISPR-Cas system has significantly advanced genome editing, offering vast applications in medical treatments and life sciences research. Despite their immense potential, the existing CRISPR-Cas proteins still face challenges concerning size, delivery efficiency, and cleavage specificity. Addressing these challenges necessitates a deeper understanding of CRISPR-Cas proteins to enhance the design and discovery of novel Cas proteins for precision gene editing. In this study, we performed extensive deep-learning research on CRISPR-Cas proteins, aiming to develop a classification model capable of distinguishing CAS from non-CAS proteins, as well as discriminating sub-categories of CAS proteins, specifically CAS9 and CAS12. We developed two types of deep learning models: 1) a transformer encoder-based classification model, trained from scratch; and 2) a large protein language model fine-tuned on ProtBert, pre-trained on more than 200 million proteins. To boost learning efficiency for the model trained from scratch, we introduced a novel margin-based loss function to maximize inter-class separability and intra-class compactness in protein sequence embedding latent space of a transformer encoder. The experimental results show that the Fine-Tuned ProtBert-based (FTPB) classification model achieved accuracies of 99.06%, 94.42%, 96.80%, 97.57% for CAS9 vs. Non-CAS, CAS12 vs. Non-CAS, CAS9 vs. CAS12, and multi-class classification of CAS9 vs. CAS12 vs. Non-CAS, respectively. The Latent Space Regularized Max-Margin Transformer (LSRMT) model achieved classification accuracies of 99.81%, 99.81%, 99.06%, 99.27% for the same tasks, respectively. These results demonstrate the effectiveness of the proposed Max-Margin-based latent space regularization in enhancing model robustness and generalization capabilities. Remarkably, the LSRMT model, even when trained on a significantly smaller dataset, outperformed the fine-tuned state-of-the-art large protein model. The high classification accuracies achieved by the LSRMT model demonstrate its proficiency in identifying discriminative features of CAS proteins, marking a significant step towards advancing our understanding of CAS protein structures in future research endeavors.

## I. Introduction

The Clustered Regularly Interspaced Short Palindromic Repeats (CRISPR) and its associated proteins (Cas) form the cornerstone of the revolutionary CRISPR-Cas system, a groundbreaking technology that has revolutionized gene editing since its inception [1]. This system, functioning as molecular scissors, enables precise gene modifications within disease genomes, thereby opening new therapeutic possibilities for a variety of diseases. Central to the CRISPR-Cas system are the Cas proteins, which, as endonucleases, work in tandem with guide RNA (gRNA) to identify target DNA sequences, a process facilitated by the Protospacer Adjacent Motif (PAM). With the Cas protein family, Cas9 (a type II endonuclease) and Cas12 (a type IV endonuclease) stand out prominently due to their extensive utilization in CRISPR-Cas gene-editing experiments [2, 3]. However, the limited availability of annotated Cas proteins poses a significant challenge in fully harnessing the CRISPR-Cas system’s potential and understanding its therapeutic capabilities. The current Cas proteins confront challenges, including low specificity, off-target effects, and inefficient delivery mechanisms. In light of these challenges, the pursuit of innovative Cas proteins is imperative to develop more precise gene-editing tools.

While many computational tools for the CRISPR-Cas system have been developed in different studies, the task of identifying novel Cas proteins within bacterial proteomes remains a formidable challenge. Existing tools, primarily designed for the detection of CRISPR arrays in bacterial genomes, often overlook the specific identification of Cas proteins themselves. Current Cas prediction tools, such as HMM-CAS [4] and CasPredict [5], relying on Hidden Markov Models and Support Vector Machines, respectively, exhibit limitations in terms of their performance and the diversity of features they can analyze. These tools often fall short of capturing the comprehensive sequential profile of protein sequences.

In response to these limitations, this study delves into the application of advanced deep learning architectures, notably transformers, known for their big success in diverse natural language processing tasks. We investigated transformer-based deep learning architectures to understand protein sequence languages and successfully developed new deep learning models to achieve precise classification of Cas9 and Cas12 proteins amidst Non-Cas proteins. This research represents a pioneering endeavor, dedicated to the classification of Cas proteins, to advance our understanding of their sequence representations. The classification models developed in this study not only serve as robust tools for identifying new Cas12 and Cas9 proteins but also possess the potential to guide the design of novel Cas proteins with improved cleavage properties to advance the field of precision gene editing.

This paper is structured as follows: Section II provides an overview of relevant prior research. Section III presents an in-depth exploration of the methodology, elucidating the proposed deep-learning models for Cas protein classification. In Section IV, we present the experimental results, conducting a comprehensive analysis of the effectiveness and efficiency of our proposed methods. Lastly, in Section V, we conclude this study and outline avenues for future research.

## II. Related Works

The integration of machine learning (ML) and deep learning (DL) techniques into CRISPR-Cas protein research represents a significant evolution in the field, transitioning from basic computational methods to advanced predictive models. This section reviews related contributions and how they paved the way for the methodologies presented in this study.

Early computational efforts were instrumental in identifying CRISPR arrays and Cas proteins, utilizing sequence alignment and pattern recognition techniques. Tools such as CRISPRFinder [6], CRISPRCasFinder [7], CRISPRDetect [8], and CRISPRCasTyper [9] laid the groundwork for cataloging CRISPR loci, although they offered limited insights into Cas protein functionality and specificity. The application of ML algorithms, including Support Vector Machines (SVM) and Random Forests (RF), were employed in different studies for the classification and prediction of Cas protein properties and cleavage efficiencies based on a broad range of features including genomics attributes and RNA thermodynamics [10]. Padilha et al. developed a CAS subtype classification tool called CRISPRcasIdentifier that applied three machine learning models SVM, Classification and Regression Trees (CART), and Extremely Randomized Trees (ERT) using hidden Markov model (HMM) extracted features of CRISPR cassettes data (genes encoding of Cas proteins) [11, 12]. Yang et al. developed an SVM model and obtained an accuracy of 84.84% for Cas vs. non-Cas protein classification using 400 dipeptide composition features extracted from protein sequences [5]. Despite their promising results, these models often required extensive feature engineering and domain-specific knowledge.

The advent of deep learning revolutionized the field of protein data modeling and analysis. The deep learning methods for sequential data modeling like Convolutional Neural Networks (CNNs) and Recurrent Neural Networks (RNNs) have shown superior capabilities to capture complex patterns in protein sequences and structures, significantly reducing the need for manual feature extraction [13, 14]. These models demonstrated superior performance in predicting protein functions and interactions. Zhang et al. developed a hybrid CNN-SVR model for on-target efficacy prediction within the CRISPR-Cas12 system [15]. Li et al. integrated CNNs with XGBoost for sgRNA on-target efficacy prediction for CRISPR-Cas12 proteins [16]. Kirillov et al. combined capsule neural networks with Gaussian Processes to gRNA cleavage efficiency for both Cas9 and Cas12 proteins [17].

Drawing inspiration from their success in NLP, transformer models are exceptionally adept at processing sequential data, which makes them ideally suited for modeling protein sequences. Transformer models have been skillfully adapted to decode the intricate language of proteins, leveraging their strengths to navigate the complexities of biological sequences. Oliveira et al. employed Transformer models for protein function annotation, showcasing their proficiency in decoding amino acid sequence patterns [18]. Hu et al. developed a graph-based Transformer model to predict protein normal mode frequencies and demonstrated the superior capabilities of transformer models to predict protein properties using protein sequence data [19]. Du et al. and Tang et al. employed transformer models to predict human secretory proteins and identify plasmid contigs, respectively, exemplifying the versatility and potential of transformer models to advance our comprehension of protein sequences and structures across diverse contexts [20, 21]. Most recently, Wan and Jiang (2023) developed TransCrispr, a hybrid Transformer and CNN model to predict cleavage efficiency for CRISPR-Cas9, and showed superior prediction accuracy and generalization ability compared to CNN and RNN-based methods. [22].

Furthermore, transformer models have revolutionized the field by enabling the development of large Protein Language Models (LPLMs), which have emerged as transformative tools in computational biology and bioinformatics [23, 24]. Similar to large Protein Language Models (LLMs) of NLP trained on large corpora of words [25], similar efforts have been applied to train LPLMs using large protein databases such as BFD100, UniRef50, and UniRef100 with trillions of protein sequences. The training of LPLMs involves leveraging large protein sequence corpora to predict masked amino acids, with the learned protein representations being utilized in downstream applications [26]. Several LPLMs have been developed in recent years, including ProtBERT [27], ESM [28], xTrimoPGLM [29], Ankh [30], RITA [31], ProtGPT2 [32]. Notably, the ProtBERT family of pre-trained models, developed by Rostlab, stands out for its popularity and active open-source maintenance. The user-friendly packages enable convenient fine-tuning for diverse downstream applications, making it a versatile tool for protein research. ProtBERT-based models have been successfully applied to various protein classification tasks and demonstrated their ability to understand nuanced sequence differences [33].

Although there have been many machine learning and deep learning studies for CRISPR-Cas system, most existing works are focused on Cas protein properties, such as genome editing efficiency and specificity prediction, editing outcome prediction, and high-activity gRNA design [34]. However, there are only a few computational methods specifically designed for CAS protein identification. Currently, only a limited number of Cas proteins have been expert-reviewed and experimentally verified in protein databases. To provide more data resources for CRISPR-Cas and precision gene editing research, there is a pressing need to develop effective computational methods to accurately identify Cas proteins and discover new Cas proteins to enrich the CRISPR-Cas family. This study aims to facilitate the discovery of new Cas proteins from unknown protein sequences from existing databases as well as massively AI-generated protein sequences. We investigated transformer-based models training from scratch and a fine-tuned large protein language model based on ProtBert. A margin-based loss function is proposed to augment learning performance for the lightweight transformer model trained from scratch. Both models achieved high prediction accuracy in distinguishing CAS and non-CAS proteins, as well as CAS subtypes. The proposed lightweight margin-regularized model achieved the best prediction performance compared to the unregularized model and the fine-tuned large protein model. In the next section, we will present the proposed methodology in detail.

## III. Methodology

### A. Dataset

To explore CRISPR-Cas proteins, we prepared a dataset 287 Cas12 collected from UniProt [35] and RCSB-PDB [36], and 837 Cas9 from InterPro database [37]. The selected Cas12 and Cas9 proteins were all rigorously verified by experts for characteristic domains of Cas9 and Cas12. Furthermore, we collected 597 Non-Cas proteins from diverse families including DNA endonucleases, proteases, exonucleases, and helicases. This dataset was used in this study to develop deep learning models for Cas and Non-Cas protein discrimination, as well as Cas subtype identification (Cas9 vs Cas12). Figure 1 summarizes this dataset, highlighting class imbalances. Notably, the smaller number of Cas12 proteins is due to a limited number of expert-verified instances in the current protein database. This imbalance was carefully considered during model training to ensure enhanced accuracy and robustness.

**Fig. 1:**
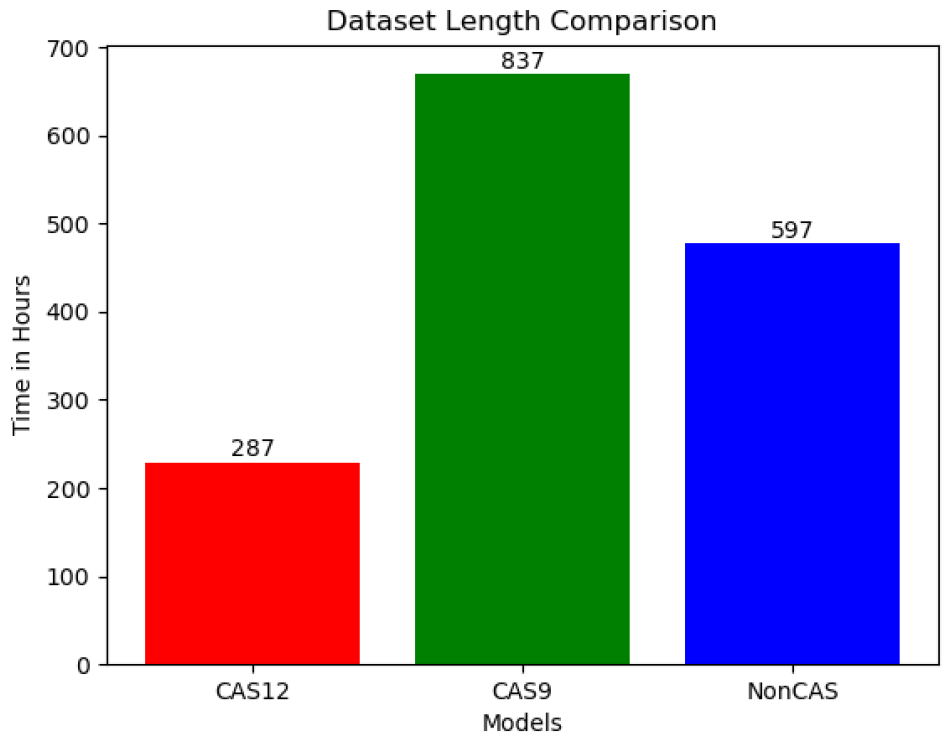
Comparison of Different Datasets we used

### B

### C. Protein Data Tokenization

We leverage the concept of proteins as life “language” and draw inspiration from cutting-edge natural language processing (NLP) techniques to unlock the secrets hidden in protein language data. Similar to treating a human language sentence in NLP, each protein sequence becomes a string of “words,” where each “word” represents an amino acid. Based on the official IUPAC one-letter notation, we define a vocabulary of 20 symbols to represent the 20 most common amino acids. Additionally, four special symbols are used to mark the beginning, end, unknown, and padding regions in a protein sequence. Just like in NLP, these symbols are known as tokens. For efficient analysis in deep learning models, each protein sequence is padded into a fixed-length sequence of 1600 tokens in our study. Then each protein sequence is transformed into a numerical representation using a predefined encoding scheme as shown in Table I. This tokenization step is crucial for standardizing protein sequences into a numerically encoded format for efficient processing by deep learning models in subsequent analytical phases.

**TABLE I:** Token Encoding for Amino Acids and Special Characters.

| Token | Value | Token | Value | Token | Value |
| --- | --- | --- | --- | --- | --- |
| A | 3 | M | 13 | V | 20 |
| C | 4 | N | 14 | W | 21 |
| D | 5 | P | 15 | Y | 22 |
| E | 6 | Q | 16 | X | 23 |
| F | 7 | R | 17 | < start > | 1 |
| G | 8 | S | 18 | < pad > | 0 |
| H | 9 | T | 19 | < end > | 2 |
| I | 10 |  |  |  |  |
| K | 11 |  |  |  |  |
| L | 12 |  |  |  |  |

### D. Transformer Based Protein Classification Model

Given the success of Transformer models in NLP, we focus on the development of Transformer models for sequential protein language data analysis. In particular, we developed a Transformer Encoder-based Protein Classification (TEP) model to discriminate different classes of protein sequences. As illustrated in Figure 3, the TEC model consists of (1) an embedding layer, followed by (2) stacked Transformer encoder layers, and (3) a final Multilayer Perceptron (MLP) based prediction head for classification tasks.

**Fig. 2:**
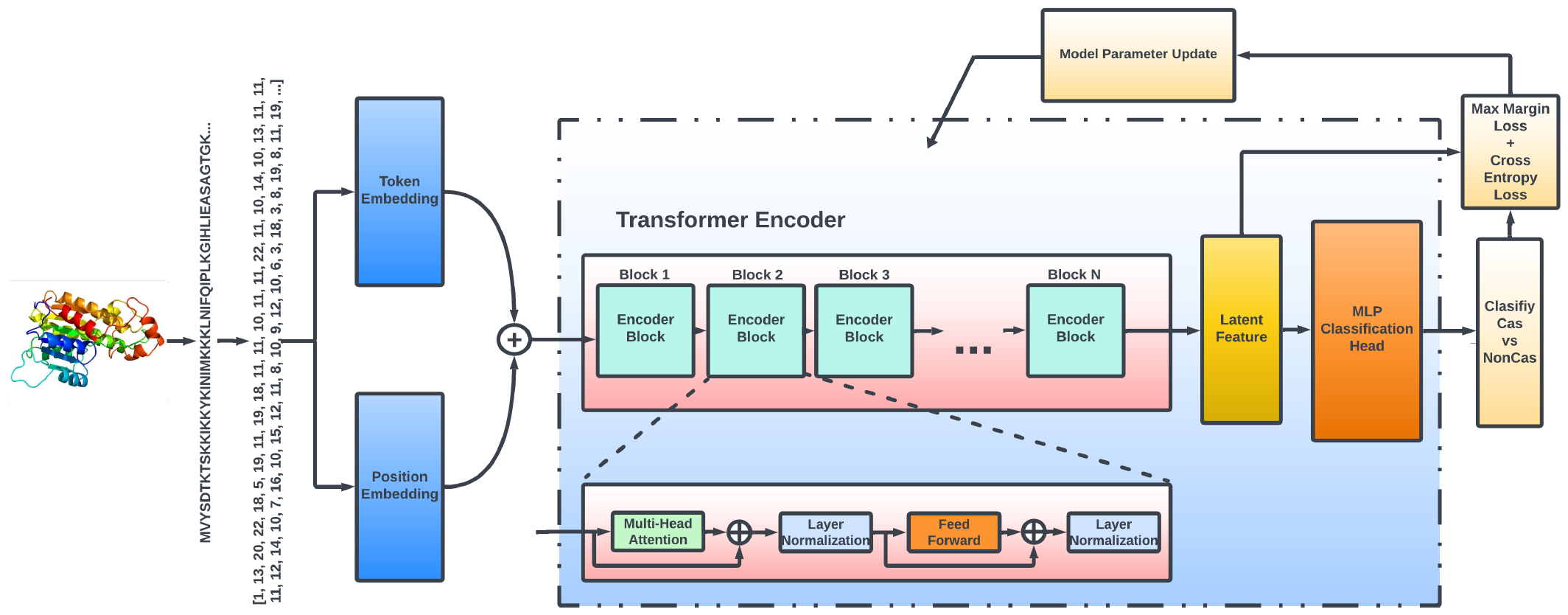
Latent Space Regularized Max-Margin Transformer Model (LSRMT)

**Fig. 3:**
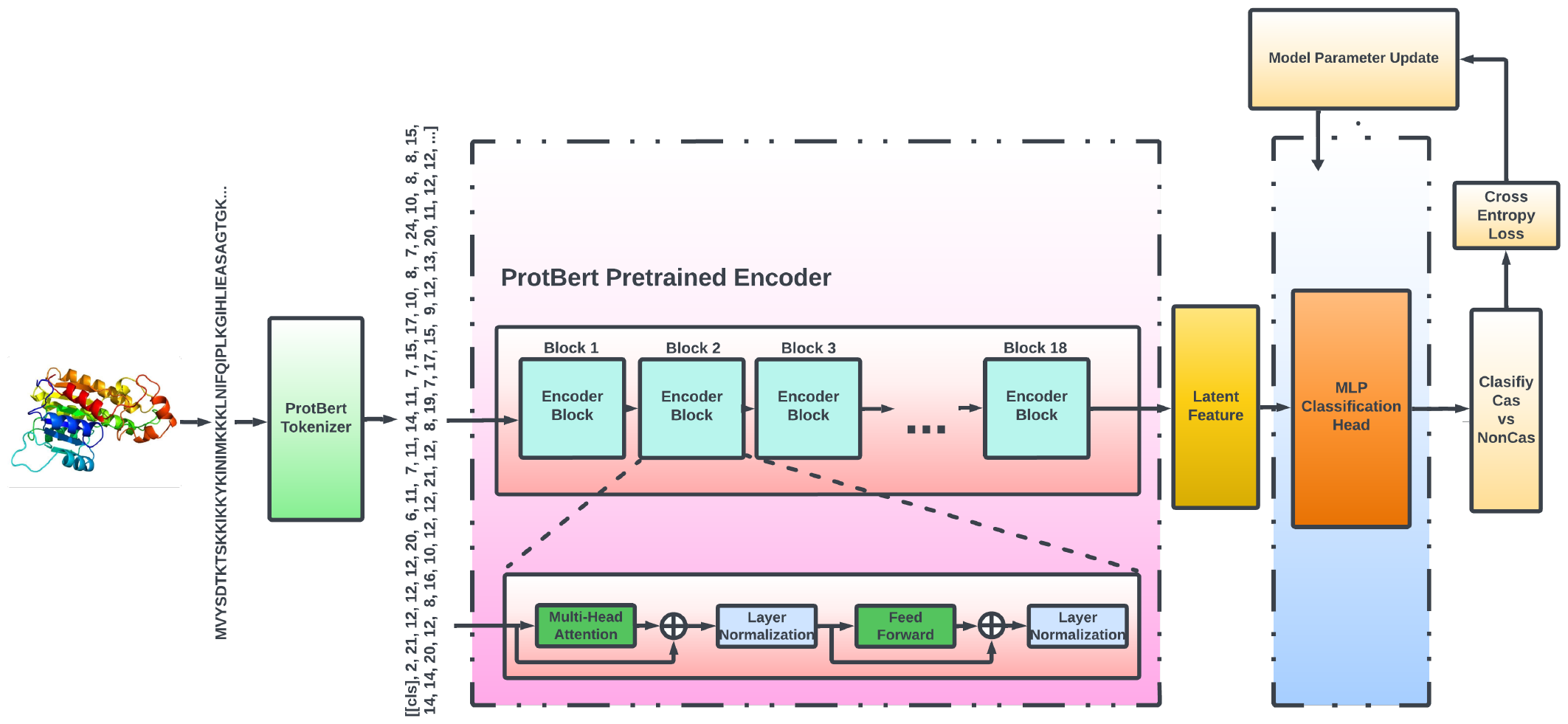
ProtBert Encoder Based Protein Classification Model (PEP)

#### Embedding Layer

the tokenized amino acid sequence padded to a fixed-length sequence of 1600 tokens is the input of the embedding Layer. We train an embedding layer that independently converts each token into a vector with dimension d = 60. The choice of 60 was made after hyperparameter optimization from d = 192, 384, 768, 60. In addition, we incorporate sequence order information by adding positional embedding to the token embedding. The final input sequence embedding is the sum of positional embedding and token embedding and is fed to the stacked transformer encoder layers for further processing. By stacking multiple Transformer encoder layers, the aim is to capture complex higher-level information and relationships from the amino acid sequence.

#### Transformer Encoder Layer

Each Transformer encoder layer contains two sub-layers – a multi-head self-attention mechanism and a fully connected feed-forward network – with residual connections around each sub-layer followed by a layer normalization operation [38]. The multi-head self-attention mechanism is composed of N separate randomly initialized attention heads. Each head is trained to identify and focus on different regions of a sequence. The self-attention mechanism is pivotal for identifying and understanding intricate relationships within protein sequences. Subsequently, a feed-forward network is used to further process the outputs of the multi-head self-attention layer to understand deeper sequence patterns and relationships. Each Transformer encoder Block also employs layer normalization and dropout strategies to enhance training stability and model generalization and mitigate overfitting issues. The Transformer encoder layer extracts the essence of a protein sequence, generating a distilled deep representation of the amino acid sequence for further analysis.

#### MLP Classification Layer

an MLP-based prediction head acts as a powerful decoder that applies a series of non-linear transformations to unlock hidden discriminative patterns for protein classifications.

### E. Max-Margin Based Latent Space Regularization

Transformer encoders excel at extracting meaningful representations that capture higher-level information and complex relationships for protein sequences. However, it faces challenges in ensuring clear separation between different protein classes in practical applications. First, the high dimensionality of extracted protein representations could make it difficult to identify crucial features to differentiate protein classes, in particular, to discriminate subtle and complicated differences of sub-types of proteins within the same family, such as the CRISPR-CAS family proteins. Also, the class imbalance can raise a potential issue that the model might prioritize the majority class in representation learning and generate poor representations for minority classes, which will increase classification errors. In addition, when trained on limited data, models can overfit the training set, performing poorly on unseen data. In particular, for those difficult data samples that are close to the decision boundaries between different classes.

In this study, we propose to introduce a Max-Margin-based latent space regularization method to address the challenges outlined above, directly targeting the issues of high dimensionality, class imbalance, and overfitting within the context of protein classification. The Max-Margin-based latent space regularization is designed to 1) maximize inter-class separability to enlarge the distance between embeddings of different protein classes in the latent space. This ensures greater distinction between protein types, leading to more accurate classifications; 2) minimize intra-class compactness to encourage embeddings within the same class to cluster closer together, promoting model confidence in recognizing similar proteins; 3) improve model’s robustness and generalization to unseen new data and potential noises in the dataset in practical applications with greater margin between different protein classes, in particular for minority classes with limited training samples.

The design of the proposed Latent Space Regularized Max-Margin Transformer Model (LSRMT) is illustrated in Figure 2. Based on the baseline structure of the TEP model as described in section III-D, we introduce a Margin loss function to enforce inter-class separability and intra-class compactness in the protein embedding latent space of the transformer encoder. Since the neural network model is updated based on mini-batch using stochastic gradient descent algorithms. The loss function is calculated based on mini-batch to update neural network weights. Given latent space feature vectors of *k* classes, each with *n*_1_, *n*_2_, *n*_3_, …, *n*_*k*_ samples respectively in a mini-batch. The Margin loss function is defined by two key terms:

1. Intra-Class Distance: the sum of the average pairwise squared distances of latent feature vectors from the same class for all *k* classes, defined by:

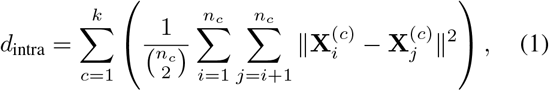

where *n*_*c*_ is the number of samples in class *c*, and 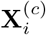 and 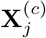 are the latent space feature vectors of the *i*th and *j*th samples in class *c*, respectively.
2. Inter-Class Distance: the sum of average pairwise squared distances between feature vectors from two different classes for all class pairs, defined by:

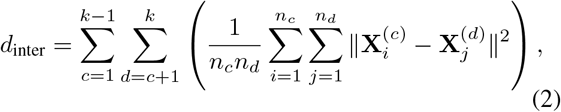

where *n*_*c*_ and *n*_*d*_ represent the number of samples in classes *c* and *d*, respectively, and 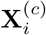 and 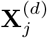 are the feature vectors of the *i*th sample in class *c* and the *j*th sample in class *d*.

The margin loss function in latent space is defined by:

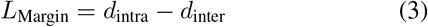

For pattern classification, a multilayer fully connected feed-forward neural network MLP is used to map the latent space feature vectors 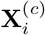 to logit vectors 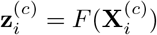, where *F* denotes the MLP neural network’s transformations. Then the softmax function is applied to these logits defined as:

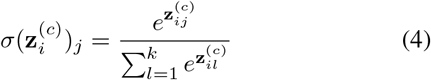

This transforms logits into class probabilities. Then the Cross-entropy loss 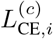 is calculated by comparing these probabilities with one-hot encoded target vectors 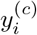. The cross-entropy loss over all samples and classes is calculated as:

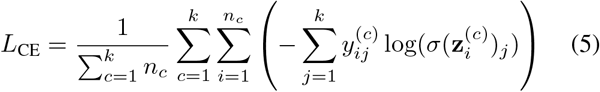

The total loss combines cross-entropy loss with margin loss from latent space vectors is defined as:

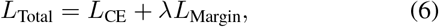

where *λ* is a regularization parameter that balances the influence of the margin loss with the cross-entropy loss. By adjusting *λ*, we can fine-tune the model’s emphasis on enhancing inter-class separability and intra-class compactness versus optimizing for classification accuracy. This allows for a flexible adaptation of the model’s learning process to prioritize feature discrimination in the latent space while maintaining high classification performance.

As illustrated in Figure 2, based on gradient descent of the loss function, the margin loss directly updates the neural network weights of the embedding layer and the transformer encoder to optimize the feature representations in the latent space for better class separability. The margin loss does not affect the classification head network that comes after the transformer encoder. In contrast, the cross-entropy loss is calculated at the last layer of the neural network. It guides the optimization to update the weights of all network layers to minimize classification errors. The adjustments made by the margin loss in the embedding and encoder layers indirectly influence the overall network performance by providing improved feature representations for the subsequent classification task handled by the classification head.

In the literature, center loss and its variants are commonly used loss functions to enhance the discrimination power of deep representation features. While these loss functions effectively promote class-center-based cohesion and separation, they may overlook local data structure nuances and neglect finer details of class distributions. To address this limitation, the proposed method leverages pairwise distances to capture intricate nuances by considering all sample interactions within a mini-batch. This approach offers a more nuanced view of class relationships, employing pairwise intra- and inter-class distance measures to enforce both intra-class cohesion and inter-class separation. While the proposed pairwise distance-based loss function carries slightly higher computational demands, the loss calculations are confined to mini-batches containing a limited number of samples (e.g., 16, 32, 64), resulting in minimal additional computational load compared to the overall neural network training process.

### F. ProtBert Based Large Protein Language Model

The field of protein sequence analysis has entered an exciting era with the emergence of Large Protein Language Models (LPLMs). These AI models, trained on massive datasets of protein sequences, are revolutionizing our understanding of protein function and interaction. ProtBert is one of the most recent cutting-edge LPLMs, which was extensively pre-trained on the Uniref100 database with 217 million protein sequences across various species [27]. The extensive pre-training process equips ProtBert with significant “linguistic” expertise, allowing it to understand the complex language of proteins and identify subtle nuances within sequences. In our study, we leveraged the power of the ProtBert for CRISPR-Cas protein research.

ProtBert is based on the Transformer architecture with an encoder block, responsible for learning protein representations, and a decoder block for protein sequence reconstruction and generation. Instead of training from scratch, we incorporated the pre-trained ProtBert encoder into our classification model architecture as shown in Figure 3. During model training, the weights of the ProtBert encoder are frozen without updating. We harnessed its pre-trained feature extraction capabilities, allowing our model to directly benefit from ProtBert’s deep understanding of protein representations. The classification head on top of the ProtBert encoder was trained specifically for our CAS protein classification tasks using cross-entropy loss. Since the ProtBert encoder cannot updated, so the margin loss is not applied in this model.

By utilizing ProtBert, we explored knowledge transfer in protein classification. We aimed to understand how pre-trained LPLMs like ProtBert can enhance protein data analysis research. This investigation demonstrates the effectiveness of LPLMs in protein sequence analysis, paving the way for advanced bioinformatics applications.

While ProtBert was powerful, we compared it to a transformer encoder-based model trained from scratch using the max-margin loss. The results, presented later, will show the superior performance of our proposed method for Cas protein classification tasks.

### G. Performance Evaluation Metrics

To comprehensively assess our models’ performance, we employed a diverse set of metrics, each offering valuable insights into different aspects of classification effectiveness. These metrics include:

**Accuracy** measures the proportion of correctly predicted instances to the total number of instances. It is a fundamental metric indicating the overall effectiveness of the model in classification tasks. While it may not fully capture the efficacy across imbalanced classes, potentially cannot reflect the model’s accuracy concerning minority classes.

**Precision** measures the model’s accuracy in predicting positive instances. It is calculated as the ratio of true positive predictions to the total number of positive predictions made. High precision indicates a low rate of false positives, which is crucial in scenarios where the cost of false positives is significant.

**Recall (Sensitivity)** assesses the model’s ability to identify all actual positive instances correctly. It is especially important in situations where missing a positive case can have serious consequences. High recall implies that the model is effective at capturing most of the positive instances.

**Specificity** measures the model’s ability to identify negative instances correctly. It is the ratio of true negative predictions to the total number of negative instances.

**F1 Score** provides a balance measure combining precision and recall with their harmonic mean. It is particularly useful in addressing imbalanced class distributions.

**Balanced Accuracy** calculates the average of Sensitivity (Recall) and Specificity, providing a holistic view considering both aspects. It is useful to provide an overall assessment of the model’s performance for imbalanced datasets, as it gives equal weight to each class.

The performance metrics are based on 4 key terms: True Positive (TP) signifies a correctly predicted positive case; True Negative (TN) indicates a correct prediction of a negative instance; False Positive (FP) an instance where the model incorrectly predicts a positive class; and False Negative (FN) happens when it misclassifies a true positive. Then the formulations of the performance metrics are defined as follows:

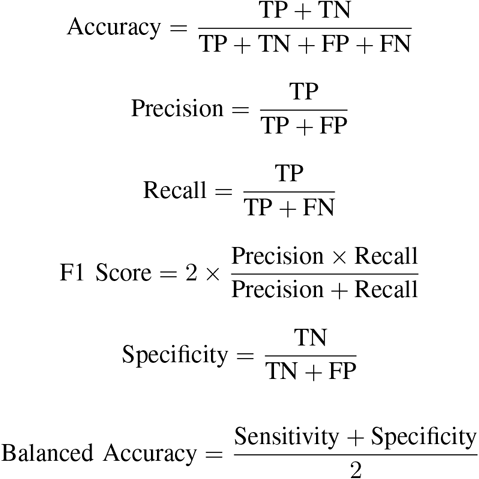

The results of our meticulous evaluation, including detailed analyses and interpretations of these metrics, are provided in the supplement section.

### H. Model Evaluation

To meticulously assess the performance of our models (TEP, LSMRT, and PEP), we employed a stringent 10-fold cross-validation approach with stratified sampling. Within each fold, we maintained the same proportion of each class as found in the overall dataset. This prevents any fold from being biased towards one class, ensuring a balanced evaluation. For each of the ten folds, we use one fold as the independent test set, and the remaining nine folds to create a training set. Furthermore, 10% of the training set is used as validation data during model training. The trained model with the best validation accuracy is selected to classify proteins in the test set. The performance metrics on the test set were recorded. This process was repeated for all ten folds. Finally, we aggregated the performance metrics across all 10 folds to obtain a comprehensive evaluation of the model’s true performance potential.

### I. Hyperparameter Optimization

Extracting the best performance from deep learning models requires exploring model structures and a lot of hyperparameters. These crucial settings control the learning process, impacting the model’s ability to identify patterns and make accurate predictions. However, with a vast array of hyperparameters and structure settings, finding the optimal combination can be a challenging task given limited computing resources. Instead of manual trial-and-error, hyperparameter optimization algorithms are powerful tools to systematically explore the hyperparameter space and identify the optimal configuration that generates the best performance. Table II summarizes the hyperparameter configurations for our model tuning, including embedding size, number of Transformer Encoder layers, attention heads, forward expansion factor, dropout rate, batch size, learning rate, and the margin loss weight *λ* in LSMRT. We employed the Tree-structured Parzen Estimator (TPE), a popular Bayesian optimization method for hyperparameter optimization [39]. Unlike simpler grid search techniques, TPE leverages a probabilistic model to predict the performance of different hyperparameter combinations. It builds a map of the hyperparameter space, where areas with higher predicted performance are prioritized for exploration, and avoid time-consuming exploration of unproductive areas. As the search progresses, TPE continuously learns and refines its map, leading it closer to the optimal configuration. By employing TPE, we were able to efficiently navigate the hyperparameter space and achieve the best performance for our models.

**TABLE II:** Hyperparameter Configurations for Model Tuning.

| Hyperparameter | Value/Range |
| --- | --- |
| Embedding Size | [10-100] |
| Number of Transformer Encoder Layers | [1-10] |
| Attention Heads | [2-16] |
| Forward Expansion Factor | [2-16] |
| Dropout Rate | [0.3-0.7] |
| Batch Size | [16,32,64,128] |
| Learning Rate | [0.0001-0.01] |
| Max-Margin Loss $\lambda$ | [0.0001-1] |

## IV. Experimental Results

Our comprehensive evaluation of three distinct deep learning models - LSRMT, TEP, and PEP - highlighted their capabilities in Cas protein classification tasks. The detailed performance metrics are summarized in Table III. The proposed LSRMT model achieved balanced accuracies of 99.58%, 97.07%, 98.37%, and 97.59% for CAS9 vs. Non-CAS, CAS12 vs. Non-CAS, CAS9 vs. CAS12, and the multi-class classification of CAS9 vs. CAS12 vs. Non-CAS, respectively. The FTPB classification model achieved balanced accuracies of 99.06%, 94.42%, 96.80%, and 88.47% for these tasks, respectively. And the non-regularized transformer encoder-based model TEP obtained balanced accuracies of 99.72%, 97.26%, 98.08%, and 98.04% for these tasks, respectively. Overall, the proposed LSRMT model achieved the best classification performance across all four classification tasks, compared to the non-regularized model version TEP and the fine-tuned large protein model PEP. The experimental results indicate that the proposed margin-based latent space regularization was effective in enhancing model learning and generalization capabilities. The dedicated lightweight model LSRMT, even trained on a significantly smaller dataset, outperformed the fine-tuned state-of-the-art large protein model. The high classification accuracies achieved by the LSRMT model demonstrate its proficiency in identifying discriminative features of CAS proteins. It paves the way for research further to advance our understanding of CAS protein in future research endeavors.

**TABLE III:** Model Performance.

| Dataset | Model | Lambda Value | Accuracy | Precision | Recall | F1 Score | Specificity | Balanced Accuracy |
| --- | --- | --- | --- | --- | --- | --- | --- | --- |
| CAS9 vs CAS12 | LSRMT | $\lambda = 0.09$ | 99.53±0.0072 | 100.00±0.00 | 98.12±0.03 | 99.03±0.0148 | 100.00±0.00 | 99.06±0.0143 |
| | TEP | $\lambda = 0.0$ | 98.91±0.0100 | 99.41±0.0176 | 96.36±0.0397 | 97.80±0.0201 | 99.79±0.0063 | 98.08±0.0191 |
| | PEP | - | 97.77±0.01 | 96.78±0.03 | 94.80±0.07 | 95.52±0.03 | 92.88±0.01 | 96.80±0.03 |
| CAS9 vs NonCAS | LSRMT | $\lambda = 0.2$ | 99.84±0.0031 | 100±0.00 | 99.62±0.0076 | 99.81±0.0038 | 100±0.00 | 99.81±0.0038 |
| | TEP | $\lambda = 0.0$ | 99.77±0.0036 | 100±0 | 99.43±0.0087 | 99.71±0.0044 | 100±0.00 | 99.72±0.0044 |
| | PEP | - | 99.16±0.007 | 99.50±0.01 | 98.48±0.01 | 98.97±0.009 | 99.63±0.008 | 99.06±0.008 |
| CAS12 vs NonCAS | LSRMT | $\lambda = 0.05$ | 99.84±0.0031 | 100.00±0.00 | 99.62±0.0076 | 99.81±0.0038 | 100.00±0.00 | 99.81±0.0038 |
| | TEP | $\lambda = 0.0$ | 97.66±0.0126 | 96.81±0.0350 | 96.14±0.0287 | 96.41±0.0187 | 98.37±0.0182 | 97.26±0.0140 |
| | PEP | - | 95.03±0.02 | 92.69±0.06 | 92.69±0.07 | 92.36±0.03 | 96.16±0.03 | 94.42±0.03 |
| CAS12 vs CAS9 vs NonCAS | LSRMT | $\lambda = 0.2$ | 99.22±0.0104 | 99.47±0.0307 | 99.24±0.0307 | 99.35±0.0232 | 99.49±0.0091 | 99.24±0.0155 |
| | TEP | $\lambda = 0.0$ | 98.04±0.0127 | 97.61±0.0270 | 97.25±0.0375 | 97.39±0.0273 | 98.99±0.0088 | 98.04±0.0127 |
| | PEP | - | 98.20±0.0199 | 97.96±0.0459 | 97.57±0.0383 | 97.71±0.0369 | 99.07±0.0138 | 97.57±0.0383 |

### A. Cas12-Cas9-NonCas Binary Classification

In the CAS9 vs CAS12 binary classification task, the LSRMT model (with *λ* = 0.09) showed exceptional performance, achieving an accuracy of 99.53%, precision of 100%, recall of 98.12%, and a specificity of 100%. The resulting F1 Score and balanced accuracy were 99.03% and 99.06%, respectively. On the other hand, the TEP model, without max-margin regularization (*λ* = 0), achieved an F1 Score and balanced accuracy of 99.79% and 98.08%, respectively; and the PEP model, fine-tuned based large protein model protBert, achieved an F1 Score and balanced accuracy of 95.52% and 96.80%, respectively. Both TEP and the PEP model performed well but were outperformed by the proposed LSRMT model, highlighting the effectiveness of the max-margin-based latent space regularization in enhancing classification accuracy and reliability. This result indicates the proposed model’s strong ability to differentiate between sub-types of Cas proteins accurately.

For the CAS9 vs Non-CAS classification task, the LSRMT model achieved the highest performance metrics, including an accuracy of 99.84%, precision and specificity at 100%, recall at 99.62%, F1 score and balanced accuracy both at 99.81%. For the CAS12 vs Non-CAS classification task, the LSRMT model significantly outperformed the other models, achieving an accuracy of 99.84%, precision and specificity both at 100%, and a recall of 99.62%. This resulted in an F1 score of 99.81% and a balanced accuracy of 99.81%. The TEP model, without any regularization, and the PEP model, despite its extensive pre-training, lagged behind in all performance metrics, particularly in F1 score and balanced accuracy, underscoring the effectiveness of the proposed LSRMT model in distinguishing CAS proteins from Non-CAS proteins with high accuracy and reliability.

Figure 4 illustrates a comparative visualization of 10-fold cross-validation balanced accuracies for the three binary classification tasks (CAS9 vs. CAS12, CAS9 vs. NonCas, and Cas12 vs. NonCas) using violin plots. These plots reveal the distribution of balanced accuracies across the cross-validation folds. Notably, the LSRMT model demonstrates a higher median value and smaller variance in balanced accuracies compared to the unregularized TEP model and the fine-tuned large protein model PEP. Both TEP and PEP model showed much wider performance distributions, and the LSRMT model is superior in achieving consistent performance across different validation folds and different tasks. This visualization indicates the effectiveness of the LSRMT model in achieving higher accuracy with greater reliability and generalization in classifying CRISPR-Cas proteins.

**Fig. 4:**
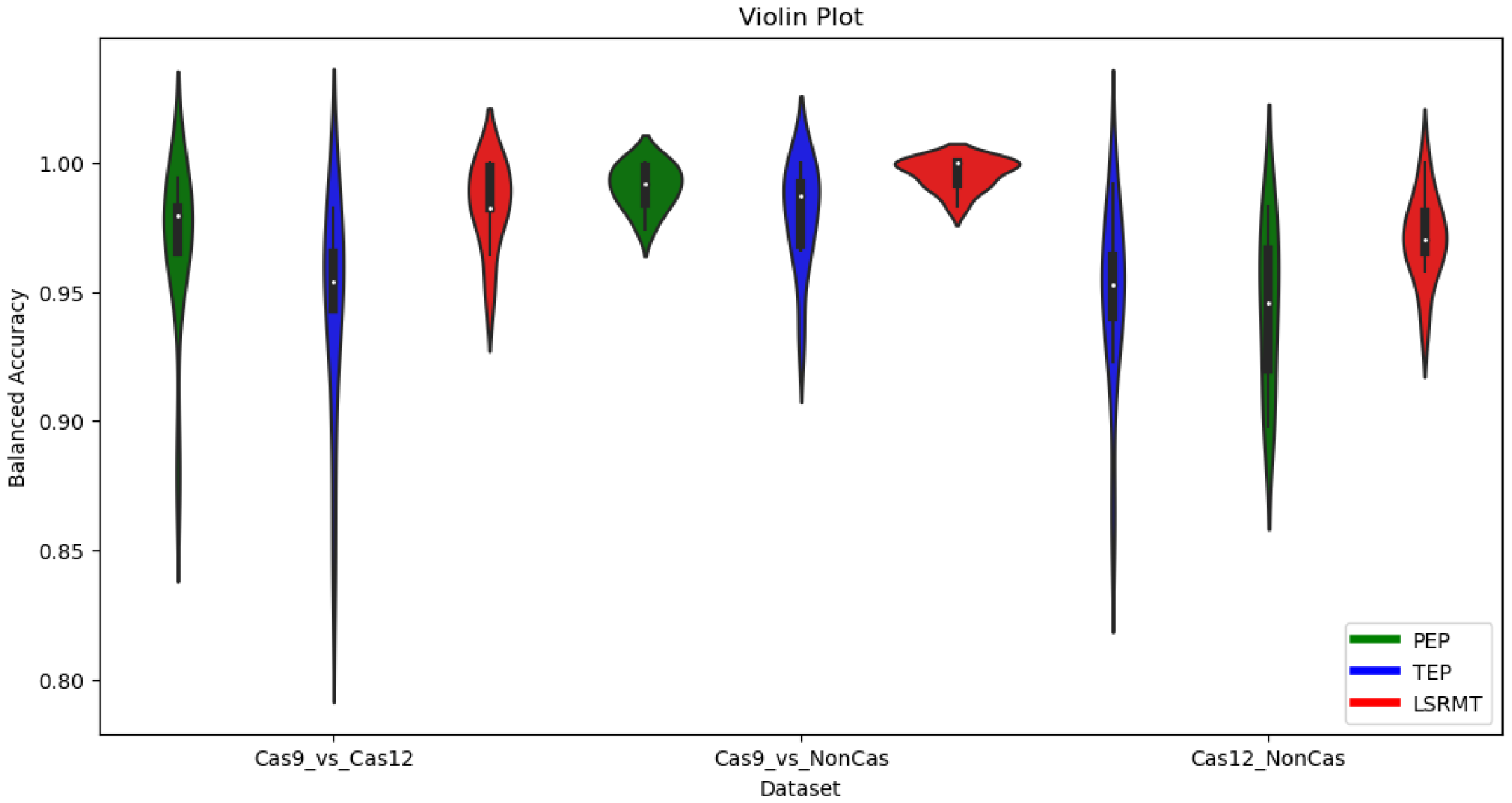
Comparative visualization of 10 fold cross validation Accuracy between ProtBert and Scratch Models.

### B. Cas12-Cas9-Non-Cas Multiclass Classification

For multi-class classification task of CAS12, CAS9, and Non-CAS proteins, the LSRMT model showed remarkable performance, achieving an accuracy of 98.59%, precision of 98.33%, recall of 98.03%, and a specificity of 99.28%. Its F1 score and balanced accuracy were 98.12% and 98.03%, respectively. These results surpassed the performance of both the TEP model and the PEP model. The superior performance of the LSRMT model in the multi-class classification task demonstrates its effectiveness in distinguishing between CAS12, CAS9, and Non-CAS proteins. Despite its lighter architecture, the LSRMT model’s exceptional performance over the fine-tuned large protein model indicates that the introduced max-margin latent space regularization that directly tackles the challenge of inter-class separability and intra-class compactness is crucial to enhancing the model’s ability to learn discriminative features for each class. On the other hand, the fine-tuning process of the PEP model, while leveraging a vast dataset, might not capture the specificities of CRISPR-Cas proteins as effectively as the model designed and trained specifically for this purpose, leading to its underperformance in this context. The proposed LSRMT’s design can be more adaptable to the specific nuances of CRISPR-Cas protein sequences, enabling more effective learning from a smaller dataset. The nuanced understanding of protein sequences through max-margin latent space regularization enables the model to accurately categorize these proteins, reflecting its potential in advancing CRISPR-Cas system research.

### C. Latent Space Analysis

Analyzing the latent space of deep learning models offers critical insights into how these models process and represent complex protein data. This analysis is vital for evaluating a model’s capability to effectively capture and differentiate between protein features. We performed a thorough latent space analysis to understand internal representations of deep learning and identify how well the models distinguish between different classes of proteins. In particular, we investigated the high-dimensional protein embedding feature vectors generated by the transformer encoder in the LSRMT and TEP models, and the ProtBert encoder for the PEP model. We applied Principal Component Analysis (PCA) to the protein embedding feature vectors and reduced the dimensionality of these embeddings to the top two principal components. This reduction enables an interpretable analysis of how each model represents and distinguishes between protein sequences in a reduced-dimensional space. The latent space visualizations in terms of the top two PCA components for the LSRMT, TEP, and PEP models across four classification tasks (CAS9 vs. CAS12, CAS9 vs. Non-Cas, CAS12 vs. Non-Cas, and multi-class CAS9 vs. CAS12 vs. Non-Cas) are depicted in Figures 5 through 8.

**Fig. 5:**
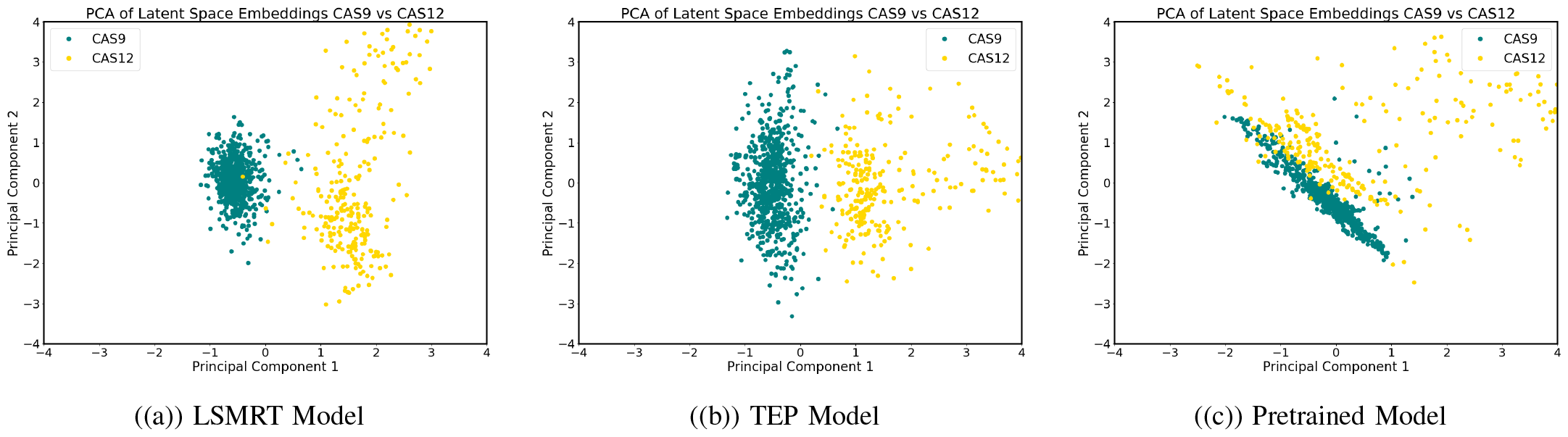
Latent Space Visualization for Dataset CAS9 vs CAS12

In Figures 5-8, PCA visualizations reveal distinct distributions and cluster formations for different classes in the latent space of the LSRMT, TEP, and PEP models. The LSRMT model, with its max-margin regularization, shows clearer class separation and denser intra-class clusters, indicating a more precise feature representation. This contrast is particularly evident when compared to TEP and PEP models, where clusters might be less distinct or more dispersed. These visualizations demonstrate the max-margin regularization’s critical role in enhancing the LSRMT model’s performance, offering a deeper insight into its advanced capability to differentiate and classify CRISPR-Cas protein sequences with high accuracy and robust generalization performance.

**Fig. 6:**
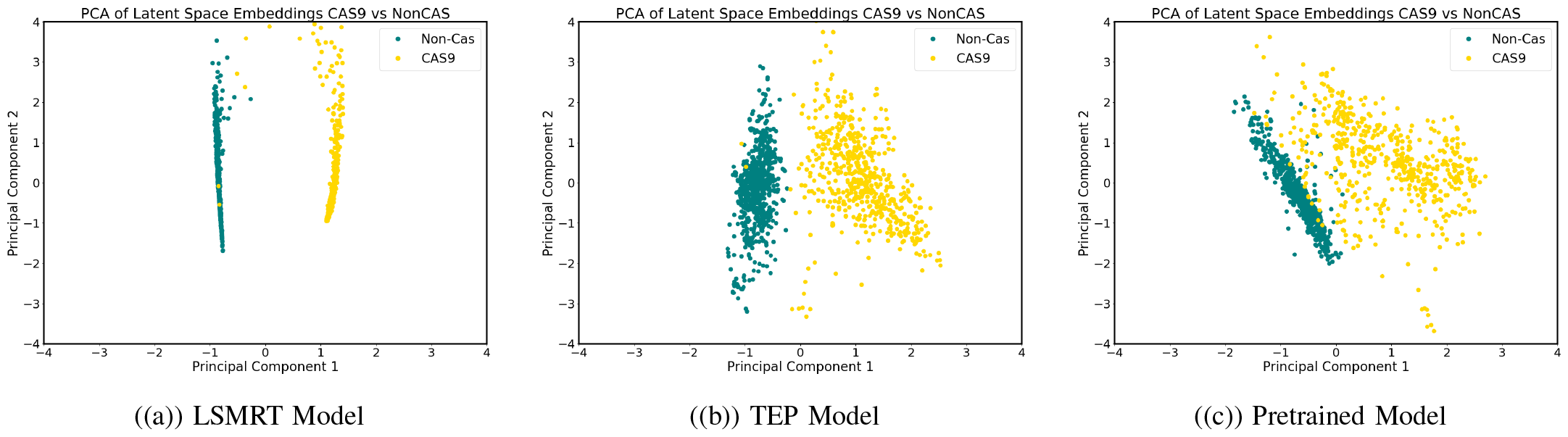
Latent Space Visualization for Dataset CAS9 vs NonCAS

**Fig. 7:**
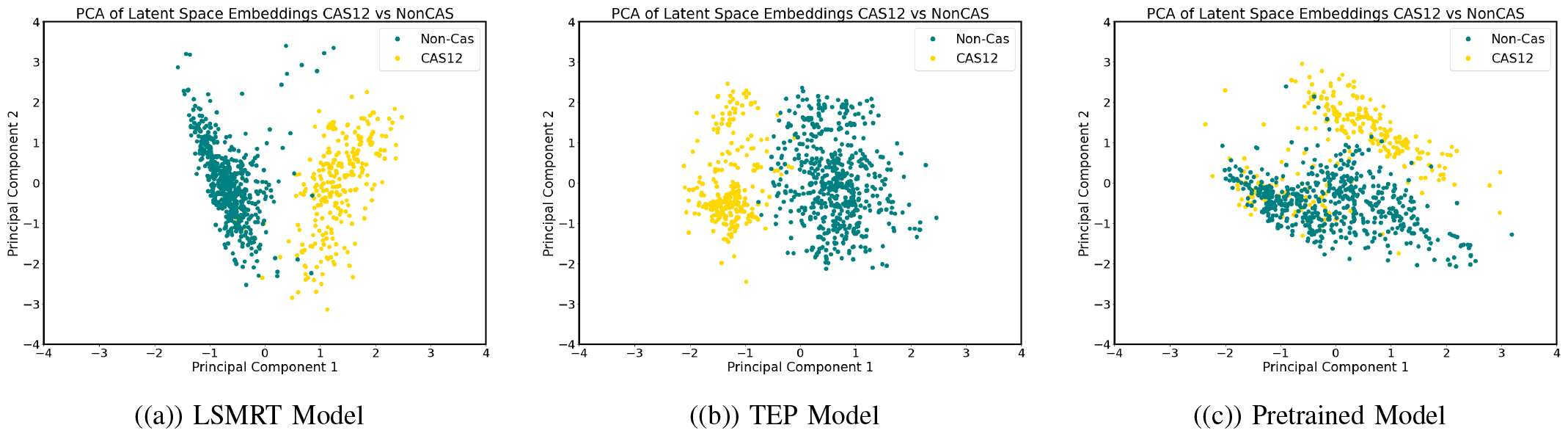
Latent Space Visualization for Dataset CAS12 vs NonCAS

**Fig. 8:**
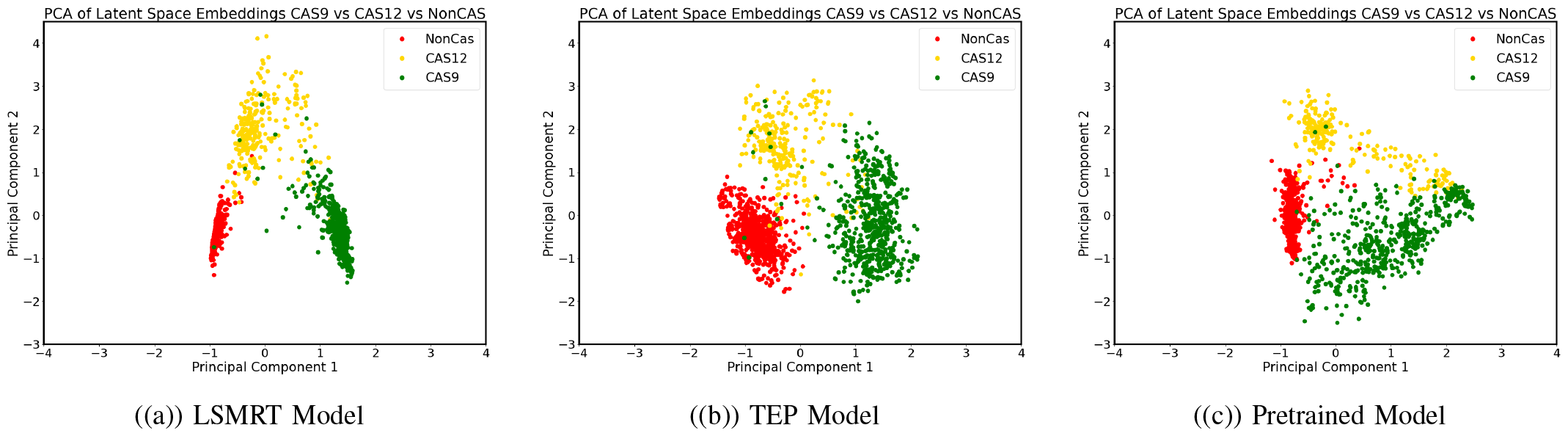
Latent Space Visualization for Dataset CAS12 vs CAS9 vs NonCAS

### D. Effectiveness of Max-Margin Regularization

To gain a deeper understanding of the impact of max-margin regularization in deep learning models, we explored the effects of varying regularization weights *λ* on classification performance. Figure 9 showed comparative plots of validation loss and accuracy during training at different *λ* values. The plots clearly demonstrate that models with max-margin regularization significantly outperform the non-regularized model. The models with non-zero *λ* values generally maintain lower validation loss and higher validation accuracy compared to models without regularization. This confirms the hypothesis that the max-margin loss function, by promoting inter-class separability and intra-class compactness, not only enhances accuracy but also reduces overfitting and enhances generalization. This analysis demonstrates the critical role of max-margin regularization in improving the generalizability and effectiveness of deep learning models in protein pattern classification tasks.

**Fig. 9:**
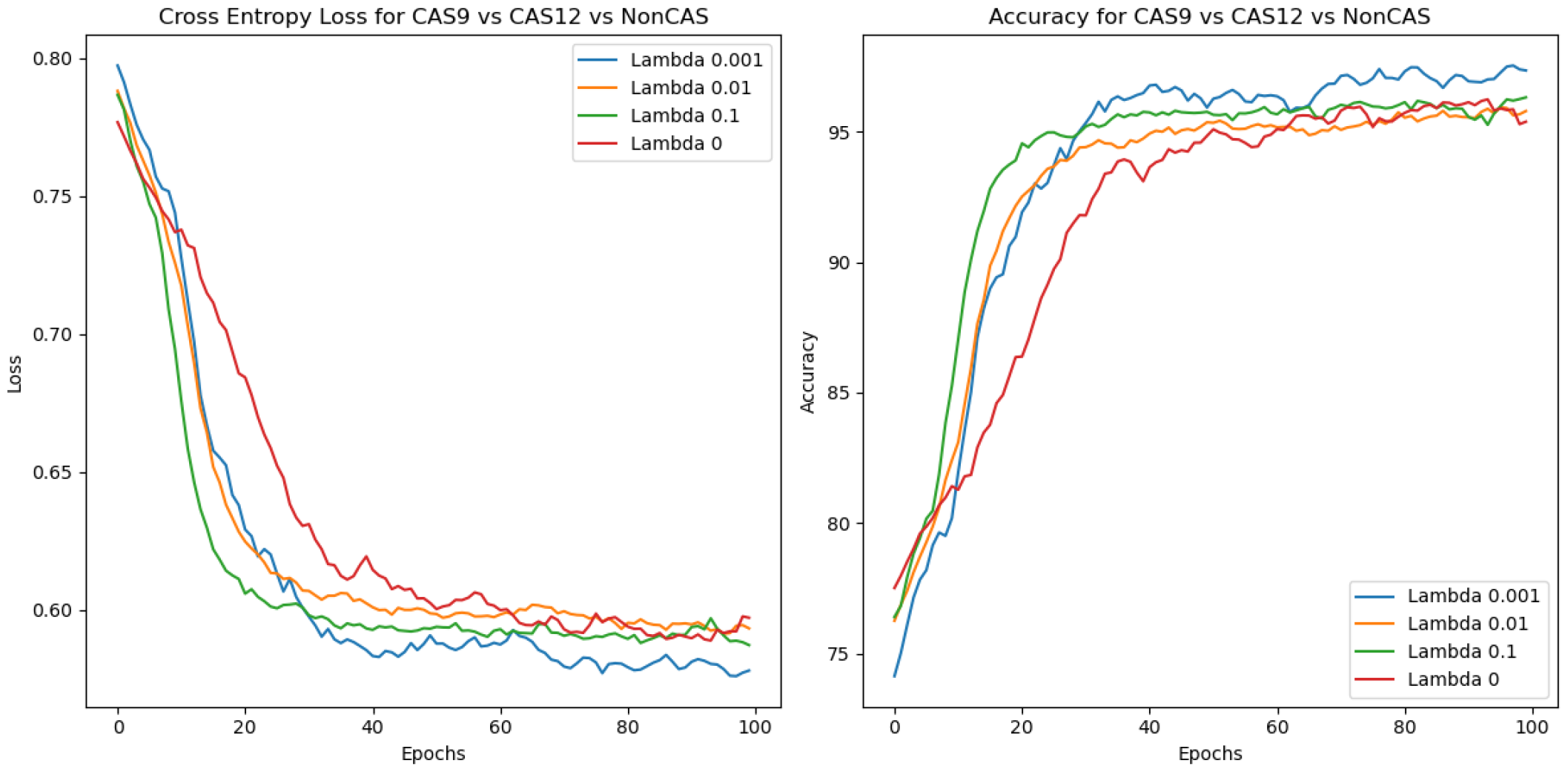
Multiclass Training Plots

### E. Computational Efficiency

As shown in Figure 10, the LSMRT model achieves quick training times without sacrificing accuracy, presenting an ideal solution for time-sensitive research. Meanwhile, the PEP model, burdened by the need to fine-tune a pre-trained network, demands greater computational resources and time, a drawback in scenarios where efficiency is crucial. This contrast highlights LSMRT’s practical advantages in delivering fast and reliable results. Moreover, it demonstrates a marginal improvement in performance, as indicated by the detailed analysis. This slight edge is attributed to the model’s innovative use of a pairwise distance-based loss function, which, despite introducing a marginally increased computational requirement, operates efficiently on small subsets of data or mini-batches. Such an approach ensures that the additional computational load remains minimal, hardly impacting the overall training process of LSMRT.

**Fig. 10:**
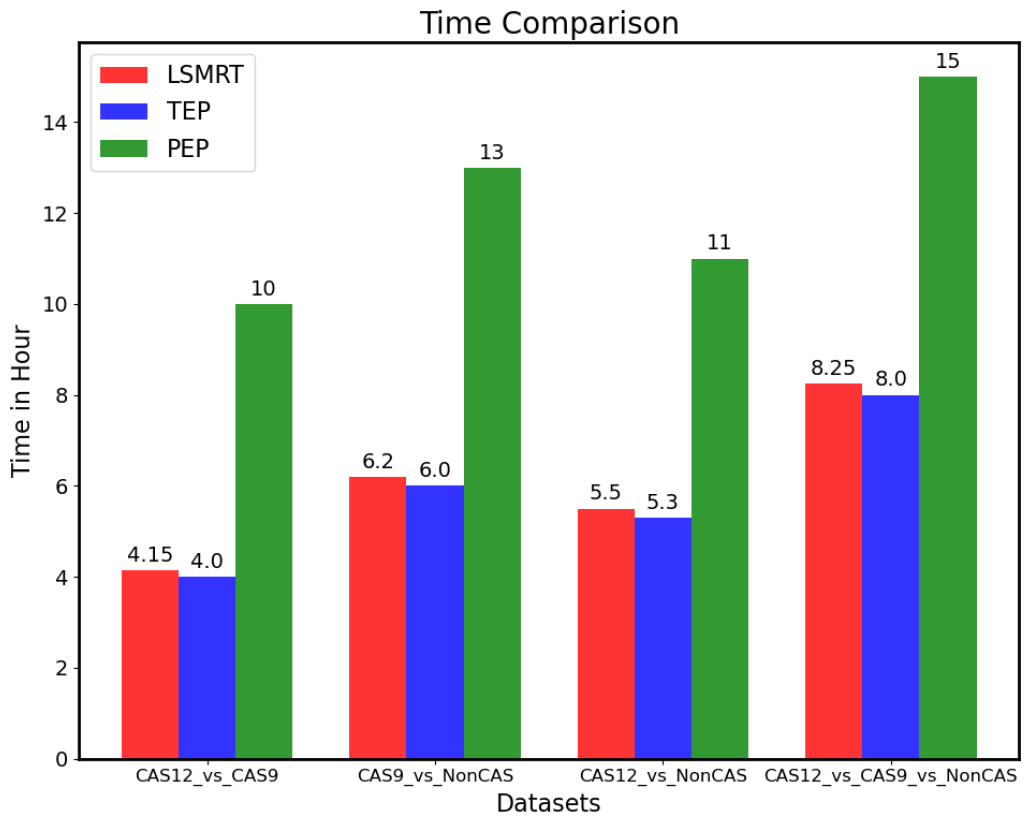
Training Time Comparison. *Note: Training performed on GPU Nvidia RTX 3090 Ti*.

### F. Future Directions in Genetic Engineering

Our study’s insights into deep learning models for protein sequence classification not only advance current methodologies but also pave the way for automation in CRISPR-CAS technology by developing state-of-the-art computational pipelines for rapid screening and characterization of novel Cas9 and Cas12 proteins, the two most widely used Cas proteins. Interestingly, the potential of these models extends far beyond conventional classification tasks, suggesting exciting possibilities for future research and practical applications. The development of hybrid models, which combine the comprehensive pre-trained knowledge base with the specificity and adaptability of scratch-trained models, stands as a promising area of exploration. Such models could offer an ideal balance between accuracy and training efficiency, essential for rapid and precise analysis in Bioinformatics. Furthermore, advancements in fine-tuning techniques could lead to highly specialized models, tailored to distinct protein sequences, thereby enhancing the precision and applicability of these models in diverse bioinformatics tasks. A particularly compelling application of these advanced models lies in the generation of efficient and smaller-size CRISPR-Cas proteins with improved editing properties. Leveraging the latent space embeddings and discriminative capabilities of models like the LSMRT [a sophisticated model excelling in both binary and multiclass protein sequence classification, leveraging latent space embeddings for enhanced discriminative capabilities, what we proposed in this paper], researchers could design and simulate novel Cas proteins with enhanced gene-editing properties. This approach could expedite the development of innovative gene-editing tools, significantly impacting therapeutic applications. Additionally, as new CRISPR-Cas proteins are discovered, the same models could be instrumental in their validation, ensuring their functionality and effectiveness in gene therapy. Further, we believe that the latent embeddings extracted from our deep learning models hold great promise for advancing generative models in AI-based protein engineering. These embeddings, rich in structural and functional relationships in proteins could guide generative models to produce novel and efficient sequences that can significantly reduce the cost associated with experimental protein synthesis. Such advancements could mark a paradigm shift in how new proteins are conceptualized, synthesized, and applied in genetic engineering. Specifically, in CRISPR-Cas technology, where generative models capable of generating novel Cas proteins are close to nil in literature, these approaches would be a guiding light for generating better and more efficient Cas proteins. In summary, the implications of our study extend beyond the current understanding of protein sequence classification. They highlight the importance of strategic model selection and application in proteomics, emphasizing performance and efficiency considerations.

## V. Conclusion

Our comprehensive investigation into deep learning for protein sequence classification has unveiled pivotal insights into the capabilities and limitations of both fine-tuned large protein models and dedicated lightweight models developed from scratch. A standout achievement of this research is the development of the LSMRT model specifically designed for protein sequence data modeling and classification. The proposed LSMRT method demonstrates superior ability in decoding complex protein sequence patterns, offering a significant improvement in both accuracy and training efficiency over the state-of-the-art large protein model fine-tuned on protbert. The LSMRT model’s agility in training and its robustness in accuracy represent a significant advancement in protein sequence classification.

Notably, the LSMRT model’s exceptional performance with over 98% accuracy in differentiating Cas from non-Cas proteins, highlights its potential as a powerful tool for identifying new CRISPR-Cas proteins within extensive databases and those generated by protein generative models. Moving forward, we are committed to further enhancing this model’s predictive accuracy and applicability. Specifically, we plan to integrate it with protein generative models to screen new Cas proteins with desirable properties, such as smaller in size, improved delivery efficiency, and cleavage specificity. This effort is directed toward advancing the discovery and design of innovative Cas proteins, thereby refining the precision of gene editing technologies in forthcoming research.

Furthermore, the LSMRT model’s quick convergence and low computational requirements highlight its suitability for rapid deployment in various research and application settings. Its deep learning architecture offers a versatile framework that is suitable for diverse sequential data modeling and classification tasks, making it an invaluable deep learning tool for the fast-paced dynamics of contemporary bioinformatics research. By setting a new benchmark for protein sequence classification, our work not only contributes to the bioinformatics field but also paves the way for future innovations in protein research and gene editing technologies.

